## Supplementary material for "Structure-based mechanism of RyR channel operation by calcium and magnesium ions": S1 and S2 Figs

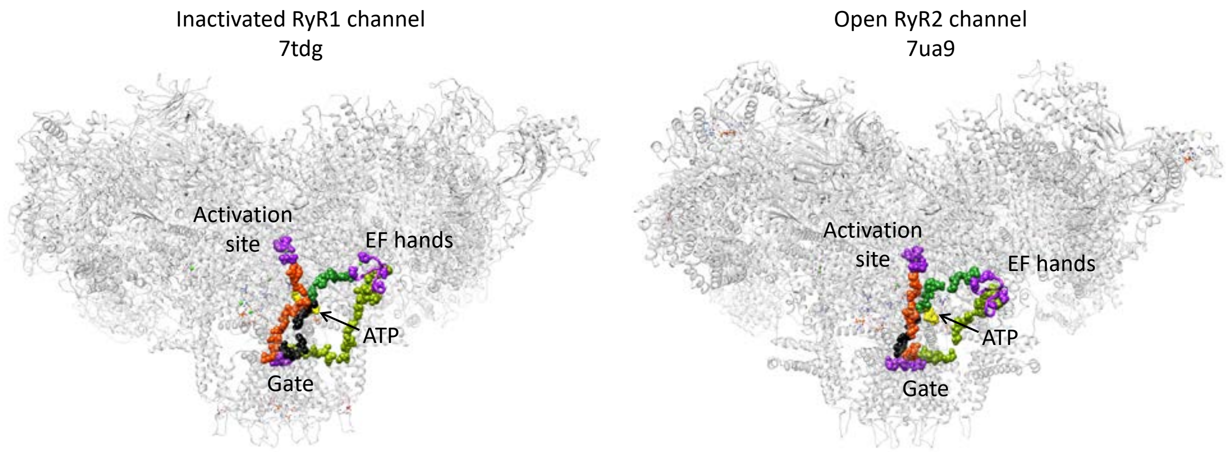


**S1 Fig. Examples of the allosteric networks in RyR1 and RyR2.** Allosteric and activation sites: purple. Activation pathways: red. Inhibition pathways: green. Common activation and inhibition pathways: black. ATP/ACP molecule: yellow.

S1 Fig complements Figure 8.

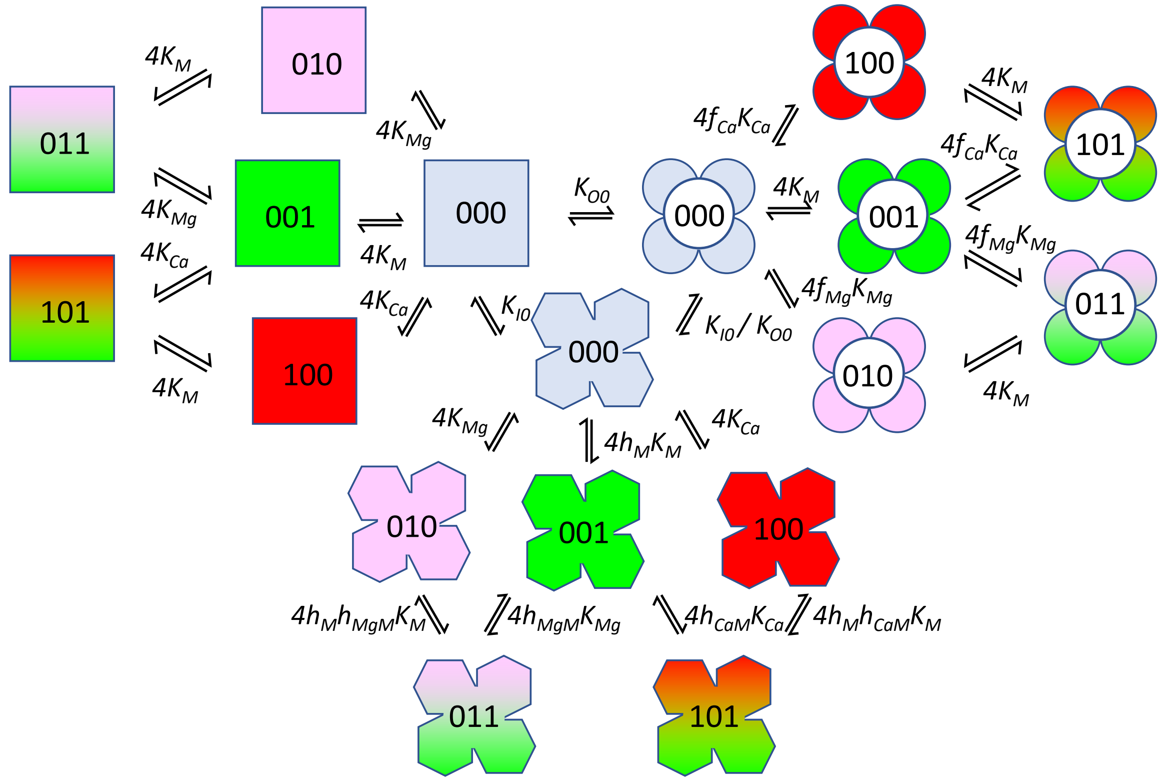


**S2 Fig. Transitions between and within macrostates of the RyR tetramer.** The overall representation of the model cannot be depicted on a two-dimensional plane, therefore the transitions between microstates are shown only for the binding of the first ion ($Ca^{2+}$ or $Mg^{2+}$) to the activation site and the first divalent ion $M^{2+}$ to the inhibition site. Other states and transitions can be constructed in the same way. The number of $Ca^{2+}$ ions bound to the activation site is i = 0 to 4; the number of $Mg^{2+}$ ions bound to the activation site is j = 0 to 4-i, and the number of $M^{2+}$ ions bound to the inhibition site is k = 0 – 4. Squares: closed state; quatrefoils with a central opening: open state; angular quatrefoils: inactivated state. The channels with 1 $Ca^{2+}$ bound to the activation site are shown in red; channels with 1 $Mg^{2+}$ bound to the activation site are shown in pink; channels with 1 $M^{2+}$ bound to the inhibition site are shown in purple. Channels with both the activation and the inhibition site occupied by one ion are shown in the red/green or pink/green gradient. The three digits inside the shapes denote respectively, from left to right, the number of $Ca^{2+}$ ions at the activation site, the number of Mg2+ ions at the activation site, and the number of $M^{2+}$ ions at the inhibition site. Note that the ion-binding dissociation constants are 4x larger than those in Figure 9 since in the ion-free tetramer, there are 4 possibilities to bind an ion to each of the binding sites, and in a tetramer with a single bound ion, there is one possibility to dissociate the ion. In addition, since all transitions are reversible, the product of each reaction cycle in the clockwise and anticlockwise directions is equal. Transitions between macrostates are shown only for the ligand-free channel but are present between all same-colored channels.

S2 Fig complements Fig 8.
