## Supplementary material for "Structure-based mechanism of RyR channel operation by calcium and magnesium ions": S1 Table

**S1 Table. Fractional occurrence of branches in the inhibition network pathways.**

| | Inhibition branches | Fraction of paths (%) | | | | | | | | |
|---|---|---|---|---|---|---|---|---|---|---|
| | | RyR1 | | | | | RyR2 | | | |
| | | C | P | O | O$^{\dagger}$ | I$^{\dagger}$ | C | C$^{\dagger}$ | O$^{\dagger}$ | O |
| | | 7k0t | 7tzc | 7m6l | 7tdh | 7tdg | 7vmm | 7ua5 | 7ua9 | 7vmp |
| | Branches of the intra-monomeric pathway | | | | | | | | | |
| I1 | INH-EF-U-K4214-S6-GATE | | 42 | 72 | 60 | | 100 | 63 | | 100 |
| I2 | INH-EF-U-T4979-S6-GATE | | | | | | | | 18 | |
| I3 | INH-EF-U-L4985, I4218-S6-GATE | | | | | | | | 74 | |
| I4 | INH-EF-U-K4211- K4821,T4822-S6-GATE | | 53 | | | 48 | | 24 | | |
| I5 | INH-EF-U-T4979-S4828, S4829-S6-GATE | | | | 27 | | | | | |
| I6 | INH-EF-U-K4211-K4821, S4828, S4829-S6-GATE | 100 | | | 4 | | | | | |
| | Subtotal | 100 | 95 | 72 | 91 | 48 | 100 | 88$^{\#}$ | 92 | 100 |
| | Branches of the inter-monomeric pathway | | | | | | | | | |
| I7 | INH-EF-E4075/R4736*-U*-S6*-GATE | | | 14 | 9 | | | | | |
| I8 | INH-EF-S4099/I4731*-U*-S6*-GATE | | | 14 | | | | | | |
| I9 | INH-EF-K4101/D4730*-U*-S6*-GATE | | 5 | | | | | | | |
| I10 | INH-EF-K4101/I4731*-U*- R4824*, S4828*, S4829*-S6*-GATE | | | | | | | | 8 | |
| I11 | INH-EF-E4075/R4736*- R4824*, S4828*, S4829*-S6*-GATE | | | | | 40 | | | | |
| I12 | INH-EF-K4101/D4730*- R4824*, S4828*, S4829*-S6*-GATE | | | | | 12 | | | | |
| | Subtotal | 0 | 5 | 28 | 9 | 52 | 0 | 0 | 8 | 0 |
| | TOTAL | 100 | 100 | 100 | 100 | 100 | 100 | 88$^{\#}$ | 100 | 100 |

INH and GATE are defined in Table 2 of the main text. The font color of a residue indicates its partaking: blue font – in the ATP binding site; brown font – in the S45 segment; green font – in the EF-hand domain; orange font – in the S23* loop.

$^{\dagger}$ marks a pair of RyR1 and RyR2 structures obtained in the same experiment.

S1 Table complements Table 4.
