## Supplementary material for "Structure-based mechanism of RyR channel operation by calcium and magnesium ions": S2 Table

**S2 Table. Fractional occurrence of branches in the activation network pathways.**

| | Activation pathways | Fraction of paths (%) | | | | | | | | |
|---|---|---|---|---|---|---|---|---|---|---|
| | | RyR1 | | | | | RyR2 | | | |
| | | C | P | O | O† | I† | C | C† | O† | O |
| | | 7k0t | 7tzc | 7m6l | 7tdh | 7tdg | 7vmm | 7ua5 | 7ua9 | 7vmp |
| | ATP-site branches | | | | | | | | | |
| A1 | ACT-CTD-U-K4214-S6-GATE | | | 100 | | 20 | 100 | 80 | | 18 |
| A2 | ACT-(CD)-CTD-U-K4957-S6-GATE | 60 | 38 | | | | | | 7 | 21 |
| A3 | ACT-CTD-U-I4218-S6-GATE | 40 | | | | | | | | |
| A4 | ACT-CTD-U-F4959-U-S6-GATE | | | | 69 | | | | | |
| A5 | ACT-CTD-W5011-U-K4957-S6-GATE | | | | | | | | | 25 |
| A6 | ACT-CTD-U-K4211-K4821,T4822-S6-GATE | | | | | 77 | | 20 | | |
| | Subtotal | 60 | 38 | 100 | 69 | 97 | 100 | 100 | 7 | 64 |
| | S45 branches | | | | | | | | | |
| A7 | ACT-CTD-U-S4828,S4829-S6-GATE | | | | 14 | | | | | |
| A8 | ACT-CTD-U-R4824,T4825,I4826-S6-GATE | | | | | | | | 93 | |
| | Subtotal | 0 | 0 | 0 | 14 | 0 | 0 | 0 | 93 | 0 |
| | CFF-site branch | | | | | | | | | |
| A9 | ACT-CTD-W5011-U-S6-GATE | | 61 | | 17 | | | | | 36 |
| | Subtotal | | 61 | | 17 | | | | | 36 |
| | TOTAL | 100 | 99 | 100 | 100 | 97 | 100 | 100 | 100 | 100 |

ACT and GATE are defined in Table 2 of the main text. The font color of a residue indicates its partaking: blue font – in the ATP binding site; brown font – in the S45 segment; cyan font – in the caffeine binding site. † marks a pair of RyR1 and RyR2 structures obtained in the same experiment.

S2 Table complements Table 5.
