## Supplementary material for "Structure-based mechanism of RyR channel operation by calcium and magnesium ions": S3 Table

S3 Table. **Interaction between the activation network and inhibition network pathways.**

| | | ACT-CTD-EF-U-K4214-S6-G | ACT-CTD-EF-U-K4957-S6-G | ACT-CTD-EF-U-I4218-S6-G | ACT-CTD-EF-U-F4959-EF-U-S6-G | ACT-CTD-W5011-EF-U-K4957-S6-G | ACT-CTD-EF-U-K4211-K4821,T4822-S6-G | ACT-CTD-EF-U-S4828,S4829-S6-G | ACT-CTD-EF-U-R4824,T4825,I4826-S6-G | ACT-CTD-W5011-EF-U-S6-G |
|---|---|---|---|---|---|---|---|---|---|---|
| | | A1 | A2 | A3 | A4 | A5 | A6 | A7 | A8 | A9 |
| I1 | INH-EF-U-K4214-S6-G | 7m6l[$]<br>7vmm[$]<br>7ua5[$]<br>7vmp[$] | 7tzc<br>7vmp | | 7tdh | 7vmp | | | | |
| I2 | INH-EF-U-T4979-S6-G | | 7ua9 | | | | | | | |
| I3 | INH-EF-U-L4985, I4218-S6-G | | 7ua9 | | | | | | | |
| I4 | INH-EF-U-K4211- K4821,T4822-S6-G | 7tdg | 7tzc | | | | 7tdg[$]<br>7ua5[$] | | | |
| I5 | INH-EF-U-T4979-S4828,S4829-S6-G | | | | 7tdh | | | | | |
| I6 | INH-EF-U-K4211-K4821,S4828,S4829-S6-G | | 7k0t | 7k0t | 7tdh | | | 7tdh[$] | | |
| I7 | INH-E4075/R4736*-U*-S6*-G* | | | | | | | | | |
| I8 | INH-S4099/I4731*-U*-S6*-G* | | | | | | | | | |
| I9 | INH-K4101/D4730*-U*-S6*-G* | | | | | | | | | |
| I10 | INH-K4101/D4730*-R4824*, S4828*, S4829*-S6*-G* | | | | | | 7tdg[#] | | | |
| I11 | INH-E4075/R4736*-R4824*,S4828*,S4829*-S6*-G* | | | | | | 7tdg[#] | | | |
| I12 | INH-K4101/I4731*-U*-R4824*,S4828*,S4829*-S6*-G* | | | | | | | | 7ua9[$#] | |

ACT, INH, and GATE are defined in Table 2 of the main text. The font color of a residue indicates its partaking: blue font – in the ATP binding site; cyan font – in the caffeine binding site; brown font – in the S45 segment; green font – in the EF-hand domain; orange font – in the S23* loop.

[#] marks the interaction of the inter-monomeric inhibition pathway with the activation pathway of the anticlockwise-neighboring monomer.

[$] marks structures where a residue of the ATP-binding site, or the S45 segment, participates in both the inhibition and activation network.

S3 Table complements Table 6.
