## Supplementary material for "Structure-based mechanism of RyR channel operation by calcium and magnesium ions": S2 file: Description of files.txt

Each directory contains data from the analysis of structures sharing the same UNIPROT Entry Identifier (the directory name corresponds to the identifier). The sequence of the UNIPROT Entry is given in FASTA format as uname.fasta, where uname is the UNIPROT Entry Identifier. The subdirectories correspond to the individual structures analyzed by the OHM server (the subdirectory name corresponds to the PDB code of the structure). 

Each subdirectory contains:
nccc-sequence.txt: The sequence of residues in the analyzed monomer model created from the pdb file nnnn.
nccc_core__dimer.pdb: The core dimer model created from the pdb file nnnn. 
Ca_nccc__paths.txt and EF12__nnnn_paths.txt: the "-paths.txt" files created by the OHM server for the activation and inhibition networks, respectively, using the core dimer model nccc_core__dimer.pdb.
Ca_nccc__net.txt and EF12__nnnn_net.txt: the "-net.txt" files created by the OHM server for the full activation and inhibition networks, respectively, using the core dimer model nccc_core__dimer.pdb.

In all files, chain A/ denotes the chain containing the analyzed allosteric sites, and chain D/ denotes the neighboring chain, located counterclockwise when the molecule is viewed from the top (cytosolic) side.
