## Supplementary material for "Structure-based mechanism of RyR channel operation by calcium and magnesium ions": Reagents and Tools Table

**Structured Methods - Reagents and Tools Table**

*Instructions: Please complete the relevant fields below, adding rows as needed. The following page provides an example of a completed table and additional instruction for entering your data in the table.*

| **Reagent/Resource** | **Reference or Source** | **Identifier or Catalog Number** |
| --- | --- | --- |
| **Experimental Models** |  |  |
| N/A |  |  |
| **Recombinant DNA** |  |  |
| N/A |  |  |
| **Antibodies** |  |  |
| N/A |  |  |
| **Oligonucleotides and other sequence-based reagents** |  |  |
| N/A |  |  |
| **Chemicals, Enzymes and other reagents** |  |  |
| N/A |  |  |
| **Software** |  |  |
| OriginPro, Version 2024. | OriginLab Corporation, Northampton, MA, USA. |  |
| UCSF Chimera Version 1.17. | <https://www.rbvi.ucsf.edu/chimera> |  |
| CLUSTAL Omega | <https://www.ebi.ac.uk/jdispatcher/msa/clustalo> |  |
| SIAS server | <http://imed.med.ucm.es/Tools/sias.html> |  |
| MODELLER | Sali and Blundell, 1993 |  |
| MODLOOP | <https://modbase.compbio.ucsf.edu/modloop/> |  |
| MIB2 server | <http://combio.life.nctu.edu.tw/MIB2/> |  |
| PDBePISA server | <https://www.ebi.ac.uk/msd-srv/prot_int/cgi-bin/piserver> |  |
| OHM server | <https://dokhlab.med.psu.edu/ohm/#/home> |  |
| **Other** |  |  |
| RCSBB PDB database | <https://www.rcsb.org/> |  |
| UNIPROT database | <https://www.uniprot.org/> |  |
